# Molecular insights into the atypical activation mechanism of GPR156 in maintaining auditory function

**DOI:** 10.1101/2024.03.07.581839

**Authors:** Xiangyu Ma, Li-Nan Chen, Menghui Liao, Liyan Zhang, Kun Xi, Jiamin Guo, Cangsong Shen, Dan-Dan Shen, Pengjun Cai, Qingya Shen, Jieyu Qi, Huibing Zhang, Shao-Kun Zang, Ying-Jun Dong, Luwei Miao, Jiao Qin, Su-Yu Ji, Yue Li, Jianfeng Liu, Chunyou Mao, Yan Zhang, Renjie Chai

## Abstract

The class C orphan GPCR GPR156, which lacks the typical extracellular region, plays a pivotal role in auditory function through G_i2/3_. Here, we demonstrate that GPR156 with high constitutive activity is essential for maintaining auditory function, and we further present two cryo-EM structures of human GPR156. The GPR156 dimer in both the apo state and G_i3_ protein-coupled state adopt a TM5/6-TM5/6 interface, indicating the high constitutive activity of GPR156 in the apo state. The C-terminus plays a dual role in promoting G protein binding within G-bound subunit while preventing the G-free subunit from binding to additional G protein. These observations explain how GPCR activity is maintained through dimerization and provide a mechanistic insight into the sustained role of GPR156 in maintaining auditory function.

## Main

GPR156 is highly expressed in auditory hair cells (HCs), and inactivation of GPR156 leads to severe hearing impairment (*1*). Variants of GPR156 identified in human pedigrees have been found to be associated with decreased expression levels, causing recessive congenital hearing loss (*2*). Moreover, GPR156 is a well-conserved cell polarity determinant, and the GPR156-Gα_i_ signaling pathway directs the orientation of mechanical sensory HCs in the mouse cochlea, mouse vestibulum, and zebrafish lateral line otolith organ. A recent study suggests the absence of G_o_ expression in auditory HCs and indicates the exclusive involvement of G_i2_ and G_i3_ in mediating the orientation impact of GPR156 on HCs (*3*). In addition, a study has shown that GPR156 is also widely distributed in the rat central nervous system, but its physiological function in the central nervous system has not been studied further (*4*).

The class C GPCRs encompass metabotropic GABAB (GABA_B_) receptor, calcium-sensitive receptor (CaSR), metabotropic glutamate (mGlu) receptors, metabotropic glycine (mGly) receptor (GPR158), taste1 receptors, and several orphan receptors (*5–7*), all of which play essential roles in intercellular communication during both physiological and pathological processes (*8, 9*). Class C GPCRs differ from other types of GPCRs in that they only function as obligatory homo- or heterodimers rather than as monomers (*10–13*). The structures of class C GPCRs solved to date all share a large and varying Venus Flytrap (VFT) extracellular domain for ligand-binding, binding of which transduces conformational changes from the VFT to the 7-transmembrane domain (TMD) (*14–16*). However, GPR156 as an orphan class C GPCR possesses a unique and extremely short N-terminal sequence (45 residues) that is not long enough to form a large extracellular region with Extracellular Loop 2 (ECL2). Additionally, GPR156 exhibits high G_i_ constitutive activity (*17*) and is highly homologous to GABA_B_, but has been ruled out as a subtype of the latter (*18*), which suggests an unconventional dimeric form and activation mechanism. Nevertheless, little is known about the mechanisms, ligands, and signaling responses associated with GPR156, and its structural organization remains unclear.

Here, our *in vivo* experiments show that GPR156 not only plays a role in the establishment of hearing, but that it also has consistent and continuous activity in order to maintain normal hearing function. To shed light on the structural basis of the sustained role of GPR156, we present the cryo-electron microscopy (cryo-EM) structures of apo GPR156 and the GPR156–G_i3_ complex at a resolution of 3.09 Å and 2.40 Å, respectively. Our results unveil a novel dimeric form and activation mechanism of class C GPCRs in which a smaller extracellular region is formed solely by ECL2 and the short N-terminus in the absence of the VFT. Furthermore, the participation of transmembrane 6 (TM6) is key to the activation of class C GPCRs (*19–21*). However, in the apo state, a previously unknown homodimer interface located between TM5 and TM6 of both GPR156 subunits was identified. In contrast to established paradigms in class C GPCRs, both subunits of GPR156 exhibited a nearly identical conformation, thus maintaining the active state. Surprisingly, upon G protein coupling, no rearrangement occurred at the dimer interface, which is unprecedented in class C GPCRs. Adding to the intrigue, in the GPR156-G_i3_ complex the C-terminus assumed a dual role. It not only participated in G protein binding within the G-bound subunit, but also simultaneously occupied the bottom of the TMD in the G-free subunit in order to impede unwanted perturbations from additional G protein binding. These observations highlight the structural and functional diversity of class C GPCRs, with potentially profound implications for advancing our comprehension of the role of GPR156 in maintaining auditory function.

### Physiological effects of the GPR156–G_i_ signaling pathway on hearing at different ages *in vivo*

Although an important role for GPR156 in hearing development has been reported (*1*), it is still unknown whether GPR156 continues to play a role after auditory maturity in view of the high G_i_ constitutive activity of GPR156 (*17*). The onset of hearing in mice does not occur until postnatal day (P)12–P14, then progresses to full maturity at P28 (*22, 23*). In order to further explore how GPR156 affects hearing at different ages, we performed *in vivo* knock-down experiments by using AAV-mediated GPR156-shRNA at three time points, including the auditory development stage (P2–P3), the mature auditory stage (P30), and the late stage of auditory maturation (P60) (Fig. 1 and fig. S1). Two AAV-mediated GPR156-shRNAs were designed for delivery into the mouse cochlea through the round window membrane (fig. S1B). Of the two, GPR156-shRNA1 was used throughout the study, and it could be successfully delivered to HCs and could knock down GPR156 by about 50% at the transcriptional level (fig. S1, B and C).

As expected, the stereocilium of HCs in the GPR156 knockdown cochlea showed improper deflection during hearing development, which was consistent with the finding in GPR156 knockout mice (*1*) (fig. S1, D and E). Of particular interest is the observation that upon maturation of the auditory faculty (P30 and P60) knockdown of GPR156 leads to severe hearing loss (Fig. 1, C and I) and partial loss of HCs (Fig. 1, D, E, G, J, K, and M). The above *in vivo* experiments demonstrate that after hearing maturity, appropriate expression of GPR156 and its constitutive activity play an important role in maintaining normal hearing function. Although the specific physiological functions of GPR156-Gα_i_ and its essential role in auditory establishment have been demonstrated (*1, 3*), crucial questions remain regarding the underlying reasons for the high constitutive activity of GPR156 and the structural mechanism by which GPR156 activation triggers downstream signals to maintain the functionality of HCs.

**Fig. 1.**
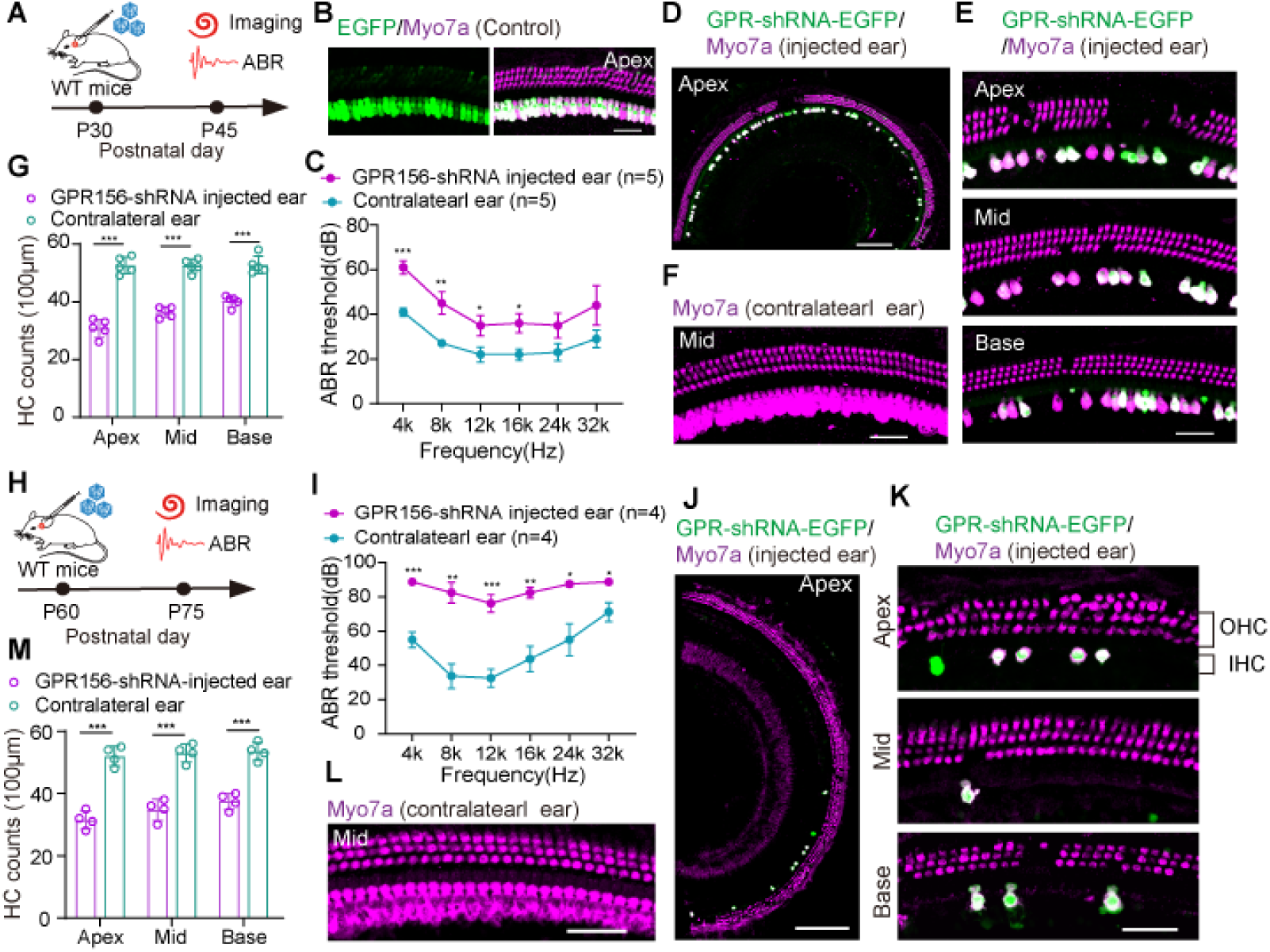
Knockdown of GPR156 causes hearing loss and hair cell loss in adult mice. **(A)** The experimental design diagram for B–G. AAV dose: 6 × 10^11^ GC /ear. **(B)** Images of the control virus infecting inner ear HCs in P30 mice. Scale bar, 40 μm. **(C)** The ABR results of the GPR156-shRNA-injected ear and the contralateral ear. **(D** and **E)** Low and high magnification confocal images of Myo7a signaling in GPR156-shRNA-injected cochleae. Apex, Mid, and Base: apical, middle, and basal turn of cochlea. Scale bar, 200 μm in D and 40 μm in E. **(F)** The confocal image of Myo7a signaling in the GPR156-shRNA contralateral cochlea. Scale bar, 40 μm. **(G)** The number of HCs in the GPR156-shRNA-injected ear and contralateral ear per 100 μm. **(H)** The experimental design diagram for I–M. AAV dose: 6 × 10^11^ GC/ear. **(I)** The ABR results of the GPR156-shRNA-injected ear and contralateral ear. **(J** and **K)** Low and high magnification confocal images of Myo7a signaling in a GPR156-shRNA-injected cochlea. Scale bar, 200 μm in J and 40 μm in K. OHC: outer hair cell. IHC: inner hair cell. **(L)** The confocal image of Myo7a signaling in a GPR156-shRNA contralateral cochlea. Scale bar, 40 μm. **(M)** The number of HCs in the GPR156-shRNA-injected ear and contralateral ear per 100 μm. Data are shown as the mean ± SEM. The *P*-value was calculated by Student’s *t-*test. \**P* < 0.05, \*\**P* < 0.01, \*\*\**P* < 0.001.

### Overall architectures of apo GPR156 and the GPR156–G_i3_ complex

As a class C GPCR, GPR156 exhibits high homology to metabotropic GABA receptors (Fig. 2A). However, GPR156 possesses a distinctively short N-terminal sequence (45 residues), which is insufficient to form a large extracellular region like typical class C GPCRs (Fig. 2, B and C). Additionally, the characteristic of GPR156’s high constitutive activity further suggests that its structure may adopt an unconventional dimeric form and activation mechanism. To determine the structural basis of the sustained role of GPR156, we expressed and purified human apo GPR156 and GPR156 in complex with the G protein G_i3_ (fig. S2 and table S1). The structure of apo GPR156 and the GPR156– G_i3_ complex were determined at global resolutions of 3.09 Å and 2.40 Å, respectively (figs. S3 and S4; and table S2).

Significantly different from other class C GPCR dimerization mechanisms, the structure of GPR156 showed a unique homodimer assembly via the ECL2, N-terminus, and TMD (Fig. 2, D and E). The conformation of the two subunits was almost the same, with the dimer showing almost mirror symmetry. High-resolution TMDs were observed for the each GPR156 protomer (fig. S5). However, N-terminus densities were limited and side chain densities were not visible due to the high conformational flexibility (figs. S3E, S4E, and S5). Nevertheless, the main organizational features of the N-terminus were identifiable. In addition, the GPR156 dimer as a whole presented a conformation in which the two subunits were close together on the intracellular side (Fig. 2, D and E). Moreover, several cholesterol molecules could be seen around the dimer (fig. S6), which is a characteristic consistent with class C GPCRs (*24*). Furthermore, cholesterol was also present within the dimer interface, while lipids packed by hydrophobic residues in the TMD could also be identified (Fig. 2, D and E, and fig. S5).

**Fig. 2.**
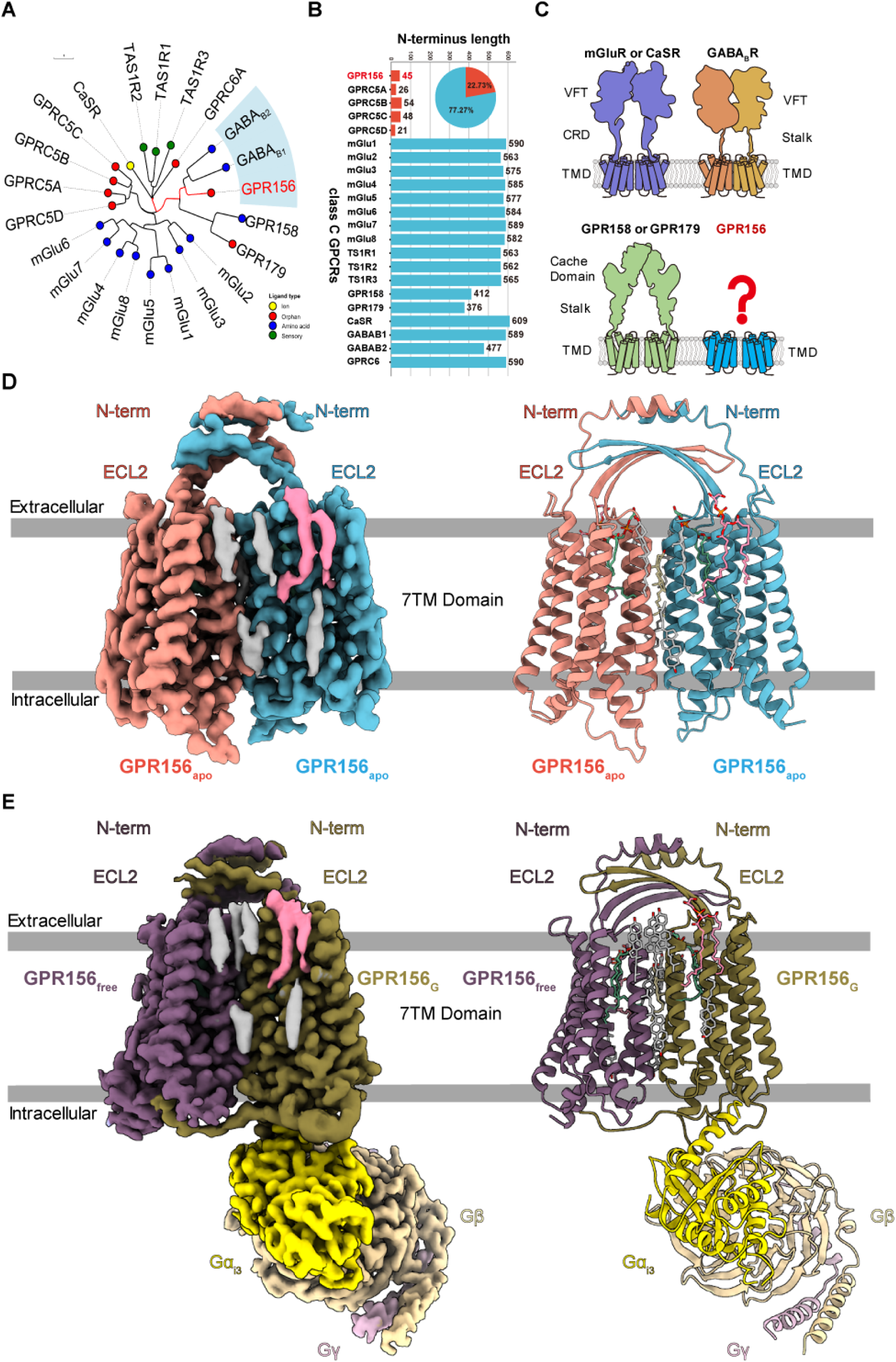
Cryo-EM structures of human apo GPR156 and in complex with Gi3. **(A)** Phylogenetic tree of the 22 class C GPCR family members with different ligand types. GPR156 is highlighted with a red mark. The scale bar indicates the number of substitutions per site. **(B)** Summary of the N-terminal lengths of the 22 class C GPCRs. GPR156 and GPRC5A–D are the only five GPCRs of class C whose N-terminal length is less than 60 residues. **(C)** The structural features of class C GPCRs. Class C GPCRs are color coded as follows. The mGluR or CaSR, GPR158 or GPR179, GPR156, GB1, and GB2 subunits are purple, green, blue, orange, and yellow, respectively. **(D** and **E)** Cryo-EM map and model of human apo GPR156 (D) and GPR156–Gi3 complex (E).

### The small extracellular region has minimal impact on the constitutive activity of GPR156

The ECL2 of the currently known class C GPCRs interacts primarily with the elongated stalk – referred to as the cysteine-rich domain (CRD) – in mGlu and CaSR (linking the VFT to the TMD) and forms a disulfide bond with the cysteine of TM3 (*20, 21, 25*). However, the N-terminus of most class C GPCRs typically comprises over 300 residues, whereas GPR156 exhibits a significantly shorter N-terminus consisting of only 45 residues (Fig. 2B). Therefore, the N-terminus of both GPR156 protomers without elongated stalks exhibit a unique conformation, forming a double-deck bridge-like structure with the ECL2 (Fig. 3A). In addition, the disulfide bond between ECL2 and TM3 (conserved C^3.29^) was previously considered conserved in class C GPCRs, but no such corresponding disulfide bond exists in GPR156 (I120^3.29^ in GPR156) (Fig. 3, B and C). Thus, the absence of this disulfide bond makes deflection of the ECL2 of GPR156 possible. This is an unprecedented configuration compared to the organization of ECL2 and the N-terminus in other class C GPCRs of various states (active, apo, and inactive states) (fig. S7, A to C).

Given the absence of the VFT region, the elongated stalk, and the disulfide bond between ECL2 and TM3, we hypothesized that GPR156’s ECL2 and N-terminus do not fulfill the conserved roles of all known class C GPCRs in response to ligand activation. To test this, we substituted the ECL2 and N-terminus of GPR156 with a linker composed of glycine and serine, respectively, and examined the effect of these on constitutive activity (fig. S7D). Consistent with our structure, these constructs exhibited slight or no alteration in basal activity when compared to wild type (WT) (Fig. 3, D and E; fig. S8A and B; and tables S3 and S4), suggesting a minimal impact of GPR156’s ECL2 and N-terminus on its constitutive activity.

**Fig. 3.**
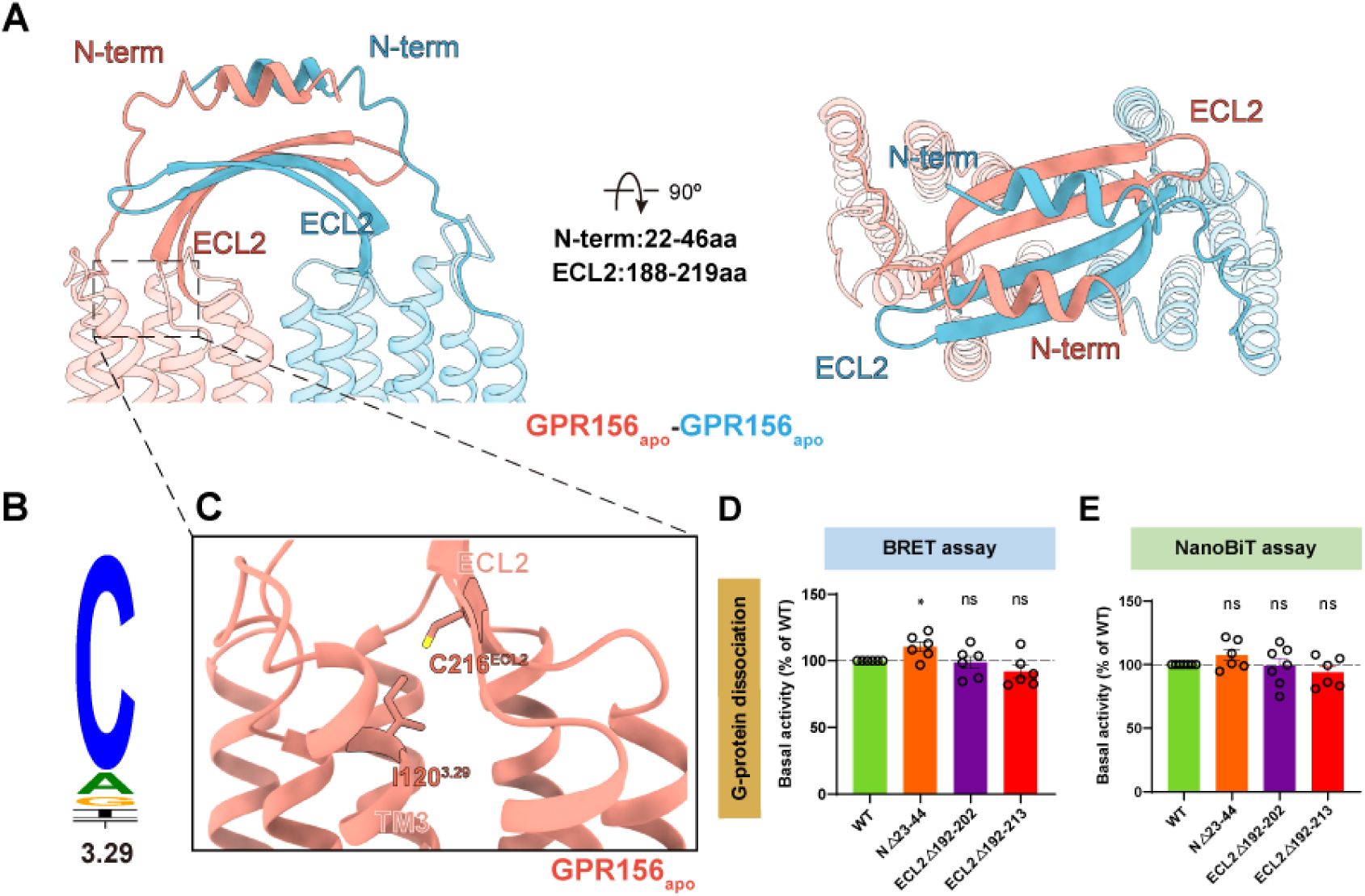
The unique arrangement of the N-terminus and ECL2. **(A)** Side view (left panel), and top view (right panel) of the distinct ECL2 and N-terminus conformations. **(B)** Sequence alignment logo showing the conservation of C^3.29^ among family C GPCRs. **(C)** Close-up view of the ECL2 and transmembrane region. In GPR156, C^3.29^ is not conservative and is replaced by I120^3.29^. **(D** and **E)** Basal activity of WT and three mutant constructs in the N-terminus and ECL2 of GPR156, measured by BRET-based assay (D) and NanoBiT-based assay (E). Data are presented as the percentage of WT activity and are shown as the mean ± SEM (bars) from at least six independent experiments performed in technical triplicate with individual data points shown (dots). \**P* < 0.05, \*\**P* < 0.01, \*\*\**P* < 0.001, \*\*\*\**P* < 0.0001 by two-tailed unpaired t-test compared to WT. Fig S8A and B provide the related surface expression level. Tables S3 and S4 provide detailed information related to BRET-based assay (D) and NanoBiT-based assay (E).

### A distinct transmembrane homodimer interface in apo GPR156

A novel unknown homodimer interface located between TM5 and TM6 of both protomers was identified in GPR156’s apo conformation (Fig. 4A, and fig. S9A). The TM5/6 helices are arranged in a V-shaped orientation at the dimeric interface (fig. S9B). In the homodimer interface, the residues at the top and bottom part of TM5 and TM6 directly contact each other to form two distinct core regions (I-II) (Fig. 4A), whereas the middle region interacts solely through a cholesterol molecule (fig. S9C). Due to the intervention of cholesterol, the area of the interaction interface increased by 165.8 Å^2^, and cholesterol has also previously been found in the middle of other class C GPCR dimer interfaces (*11, 25, 26*) (fig. S9, D and E).

The core region I, including parts Ia and Ib, encompasses the interface at the extracellular end. Ia represents a network of electrostatic interactions (D222^5.37^ and R279^6.57^; superscript numbers refer to the GPCRdb numbering scheme) that effectively tether the extracellular ends of both of the GPR156 monomers’ TMDs (Fig. 4B). Residue D222^5.37^ of each subunit also forms a hydrogen bonding interaction with Y280^6.58^ of the other protomer’s TM6, while V223^5.38^ of each subunit engages in a hydrophobic interaction with V276^6.54^ of the other subunit, both of which together establish the affiliated site Ib (Fig. 4B). The core region II covers the intracellular end of the transmembrane homodimer interface (Fig. 4C). A hydrophobic contact network is formed by L237^5.52^, V264^6.42^, and V268^6.46^ of each subunit (IIa in Fig. 4C). In addition, L237^5.52^, Y241^5.56^, and L234^5.49^ on TM5 of one protomer in this region form van der Waals forces with M261^6.39^ and N265^6.43^ on TM6 of the other protomer, respectively (IIb in Fig. 4C).

While the involvement of TM5 in the class C GPCR apo state homo- or heterodimer interface has been previously identified (apo GABA_B_ receptor (*27*) and apo GPR158 (*25, 28*)) (Fig. 4D), TM6 has previously only been shown to participate in the active class C GPCR dimer interface (*19–21*). Of particular interest, this is the first time that TM6 has also been found to be involved in an apo state dimer interface. The dimerization of GPR156 in the apo state leads to the formation of a TM5/TM6 interface, which may induce steric hindrance between the two subunits’ TM5 helices, thereby facilitating the movement of TM5 and TM3 akin to the positive allosteric modulator (PAM) function observed in the GABA_B_ receptor (fig. S9F). To determine the importance of this specific dimer interface, we mutated the dimer interface and performed BRET-based G-protein dissociation assays (Fig. 4E; fig. S8C; and table S3). It is noteworthy that the mutation to alanine in the dimer interface except R279^6.57^ and Y280^6.58^ on GPR156 reduced the basal activity of the receptor, which indicated that the TM5/TM6 interface does indeed play a role in the signal transduction of GPR156.

**Fig. 4.**
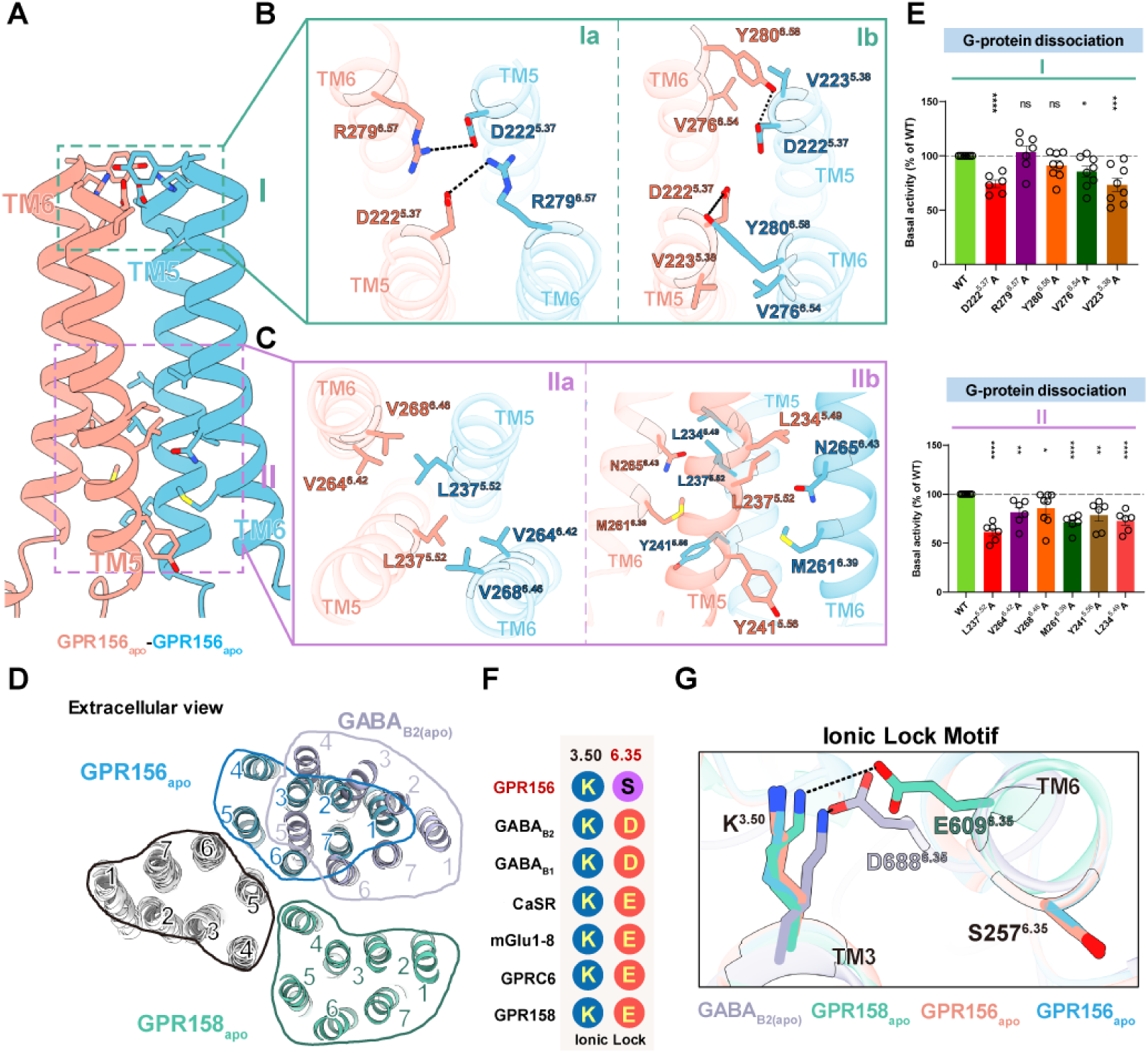
Transmembrane interface of the apo GPR156 homodimer. **(A)** Detailed interactions of the transmembrane interface in the apo state. The 7TM homodimer interface is formed at two regions (I and II), namely the extracellular and intracellular side. **(B)** Detailed interactions on the surface of region I. The network of electrostatic interactions is shown in Ia, and the affiliated hydrophobic interaction is shown in Ib. **(C)** Detailed interactions in the intracellular region II. A hydrophobic contact network is shown in IIa. The van der Waals forces at the homodimer interface are shown in IIb. **(D)** Comparison of the dimeric interface of apo GPR156 transmembrane domains with those of apo GABAB receptor (PDB 6VJM) and apo GPR158 (PDB 7EWL). **(E)** The basal activity of WT and versions with mutations in the dimer interface of GPR156, as measured by BRET-based assay. Data are presented as a percentage of WT activity and are shown as the mean ± SEM (bars) from at least six independent experiments performed in technical triplicate with individual data points shown (dots). \**P* < 0.05, \*\**P* < 0.01, \*\*\**P* < 0.001, \*\*\*\**P* < 0.0001 by two-tailed unpaired t-test compared to WT. Fig S8C provides the related surface expression level, and Table S3 provides detailed information. **(F)** The key residues in the ionic lock motif (3.50 and 6.35) are aligned among members of class C GPCRs. **(G)** Close-up view of the conserved ionic lock motif showing the different conformations between GPR156 and other members of the class C subfamily in the apo state, including GABAB receptor (PDB 6VJM) and GPR158 (PDB 7EWL).

### Non-canonical features of the apo GPR156 subunits

Conventionally, the ionic lock motif is conserved in class C GPCRs, but the GPR156 subunit exhibits peculiar differences. The ionic lock is primarily formed by the conserved K^3.50^ with D^6.35^ or E^6.35^, similar to K665^3.50^–E770^6.35^ in mGlu5 (*29*), K574^3.50^–D688^6.35^ in GABA_B2_ (*26, 30*), and K502^3.50^–E609^6.35^ in GPR158 (*25, 28*) (Fig. 4F). In GPR156, S257^6.35^ replaces the D^6.35^ or E^6.35^ that is found in other class C GPCRs, thereby potentially preventing the formation of a stable interaction with K^3.50^ in the apo or inactive state (Fig. 4, F and G).

Given the lack of the ionic lock motif and the high homology between GPR156 and the GABA_B_ receptor, comparing the TMDs of the two subunits in apo GPR156 with the GABA_B2_ subunit binding the G protein (GABA_B2(G)_) revealed a high degree of similarity, with the root mean square deviation (RMSD) measuring 1.023 Å and 0.964 Å (Fig. 5A, and fig. S10, A and B). K141^3.50^ in both subunits of apo GPR156 forms hydrogen bond with N88^2.39^, which mirrors the previously observed interaction in the GABA_B2(G)_ subunit (K574^3.50^–N520^2.39^) (*19*) (Fig. 5B). In addition, S84^2.35^ and R144^3.53^ also form hydrogen bond to further stabilize the conformation, and this previously undiscovered interaction similarly exists in the GABA_B2(G)_ subunit (Fig. 5B). The conformations of the active state feature F^3.44^ in both apo GPR156 subunits, and these closely resemble the conformation in GABA_B2(G)_, showing marked distinctions from the inactive state of GABA_B2_ (GABA_B2(inactive)_) (Fig. 5C). Coinciding with our structure, the mutation of K141^3.50^, R144^3.53^, S84^2.35^, and F135^3.44^ had a substantial impact on the basal activity (Fig. 5D; fig. S8D; and table S3). Furthermore, the forementioned findings underscore that both subunits in apo GPR156 exhibit active-state conformations and interactions consistent with the GABA_B2(G)_ subunit, and these are different from the inactive state of GABA_B2_ (Fig. 5, B and C, and fig. S10C), implying the ability of both GPR156 subunits to engage with G proteins in the apo state.

**Fig. 5.**
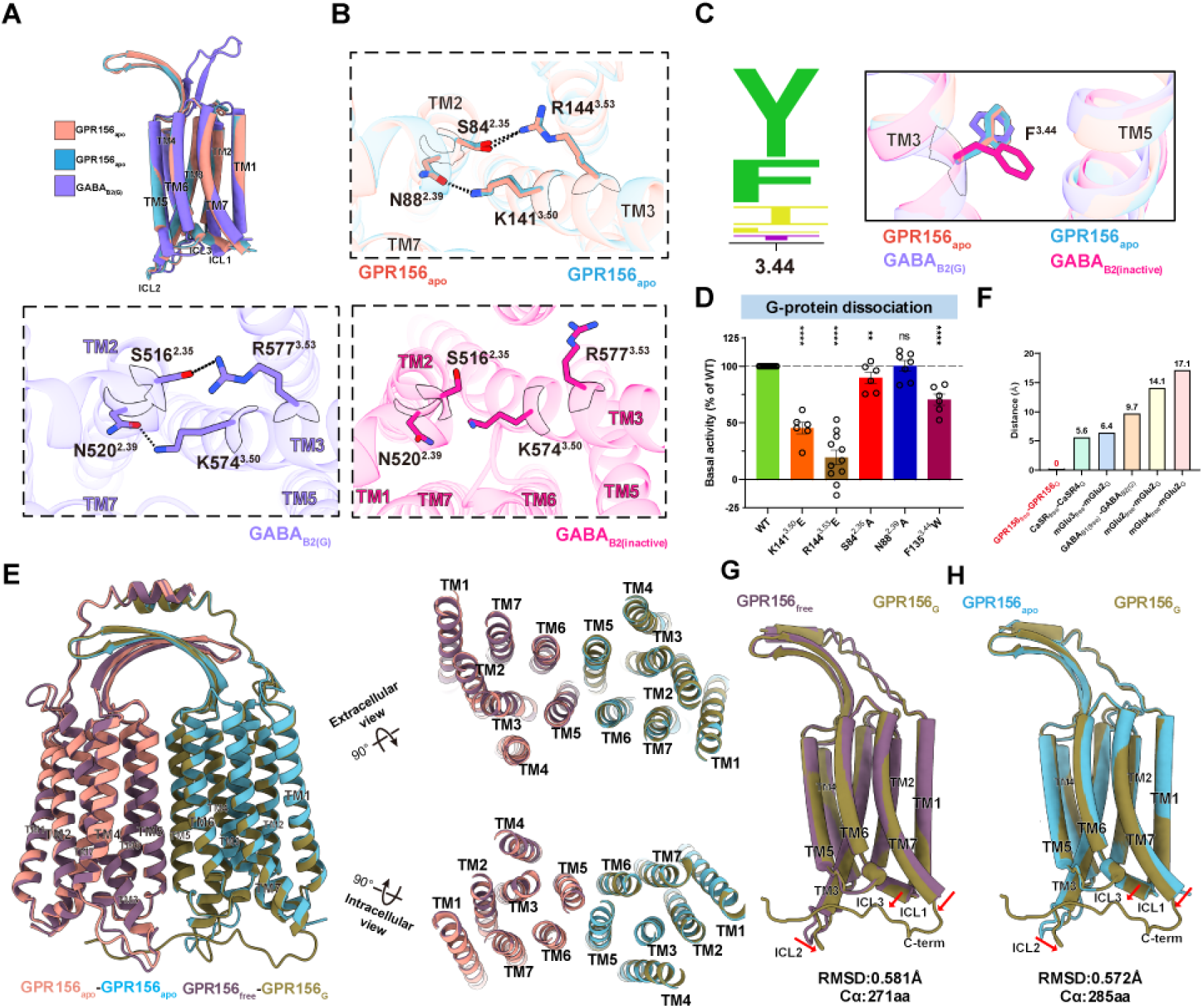
Structural details and Gi-associated transitions of the TMD in GPR156. **(A)** Side views of the superposed structures of the GPR156apo and GABAB2(G) (PDB code: 7EB2) aligned by the TMD of GABAB2(G). **(B)** Magnified views of the detailed interactions within the TMD of apo GPR156 (top panel), GABAB2(G) (left panel, PDB code: 7EB2), and GABAB2(inactive) (right panel, PDB code: 6WIV). **(C)** Sequence alignment logo showing the conservation of Y^3.29^ or F^3.29^ among family C GPCRs (left panel). Magnified views of the critical residue F in the apo state of GPR156 and different states of GABAB2 (right panel). **(D)** The basal activity of WT and mutant versions as measured by BRET-based assay. Data are presented as the percentage of WT activity and are shown as the mean ± SEM (bars) from at least six independent experiments performed in technical triplicate with individual data points shown (dots). \**P* < 0.05, \*\**P* < 0.01, \*\*\**P* < 0.001, \*\*\*\**P* < 0.0001 by two-tailed unpaired t-test compared to WT. Fig S8D provides the related surface expression level. Table S3 provides detailed information. **(E)** Side, extracellular, and intracellular views of the superposed structures of apo GPR156 and Gi-bound GRP156. Superposed structures of single protomers of apo GPR156 and Gi-bound GRP156. **(F)** Upon G protein coupling, the dimeric TMD of five class C GPCRs (including mGlu3free–mGlu2G (PDB code: 8JD3), mGlu4free– mGlu2G (PDB code: 8JD5), mGlu2free–mGlu2G (PDB code: 7MTS), CaSRfree–CaSRG (PDB code: 8WPU), and GABAB1(free)–GABAB2(G) (PDB code: 7EB2)) undergo conformational rearrangements, except for GPR156free–GPR156G. **(G** and **H)** Comparison of the GPR156 G-bound subunit and GPR156 G-free subunit (G) and the GPR156 G-bound subunit and apo GPR156 (H). The RMSD levels were calculated.

### No rearrangement occurred in the dimer interface after G protein coupling

To assess changes in GPR156 dimerization after G protein coupling, a comparative analysis between apo GPR156 and the active GPR156 dimer revealed striking similarity (RMSD of 0.494 Å) (Fig. 5E). Surprisingly, after G protein coupling, no interface rearrangement occurred in GPR156, a phenomenon unprecedented in class C GPCRs (Fig. 5, E and F). When comparing the G-bound subunit with both the G-free and apo subunits (RMSD of 0.581 Å and RMSD of 0.572 Å, respectively), we observed only minor intracellular contraction in ICL2, ICL1, and TM7 of the G-bound subunit (Fig. 5, G and H). Notably, the G-bound subunit exhibited none of the common rotations or translations found in class C GPCRs but showed the presence of the C-terminus (Fig. 5, G and H, and fig. S5). The subsequent conformation and interactions of the active state GPR156 subunits’ internal TMs mirrored the apo state conformation and interactions of both subunits in the apo GPR156, akin to the GABA_B2(G)_ subunit (Fig. 5H, figs. S10, D to F, and S11). It can be inferred that a symmetric dimeric form exists in GPR156 (apo state) with both subunits adopting an active-like state. Upon G protein coupling, no substantial rearrangements occur, with only minor intracellular region contraction in the G-bound subunit to accommodate the G protein (Fig. 5, G and H). Of particular interest, this is the first time a class C GPCR has been shown to have a dimeric interface in the active state that exists in a completely symmetric form, and this is also the first structure to show the involvement of TM5 in the active state (fig. S12).

Our structure also revealed a kind of endogenous phospholipid within the transmembrane domains of both subunits in apo GPR156 and the GPR156–G_i3_ complex structures, also located at the extrahelical site. From the density and pocket size we inferred that this was a phospholipid with two aliphatic chains, and mass spectrometry was used to further identify this as phosphatidylglycerol (PG) 36:2 (fig. S13, A to D, and data S1). It is noteworthy that each GPR156 subunit establishes a comprehensive network of contacts with both bound lipids, and this may help maintain the stability of the receptor (fig. S13, E and F). In addition, three replicates of the GPR156 dimer system with the internal phospholipids removed were further created, and a 300 ns molecular dynamics (MD) simulation was sampled for each replicate (fig. S13, G and H, and data S2). Based on the calculation results of the cavity volume for all three MD trajectories, a similar collapse of the transmembrane helices was observed, as demonstrated in the GABA_B_ receptor (*31*). Like the GABA_B_ receptor (C^6.50^, corresponding to W^6.50^ of the toggle switch motif) (*26, 30*), G^6.50^ of GPR156 lacks a large side chain (fig. S13, I to K), preventing it from responding to the activation of an intramembrane ligand as observed in mGlu2 (*32*) (fig. S13L).

**Fig. 6.**
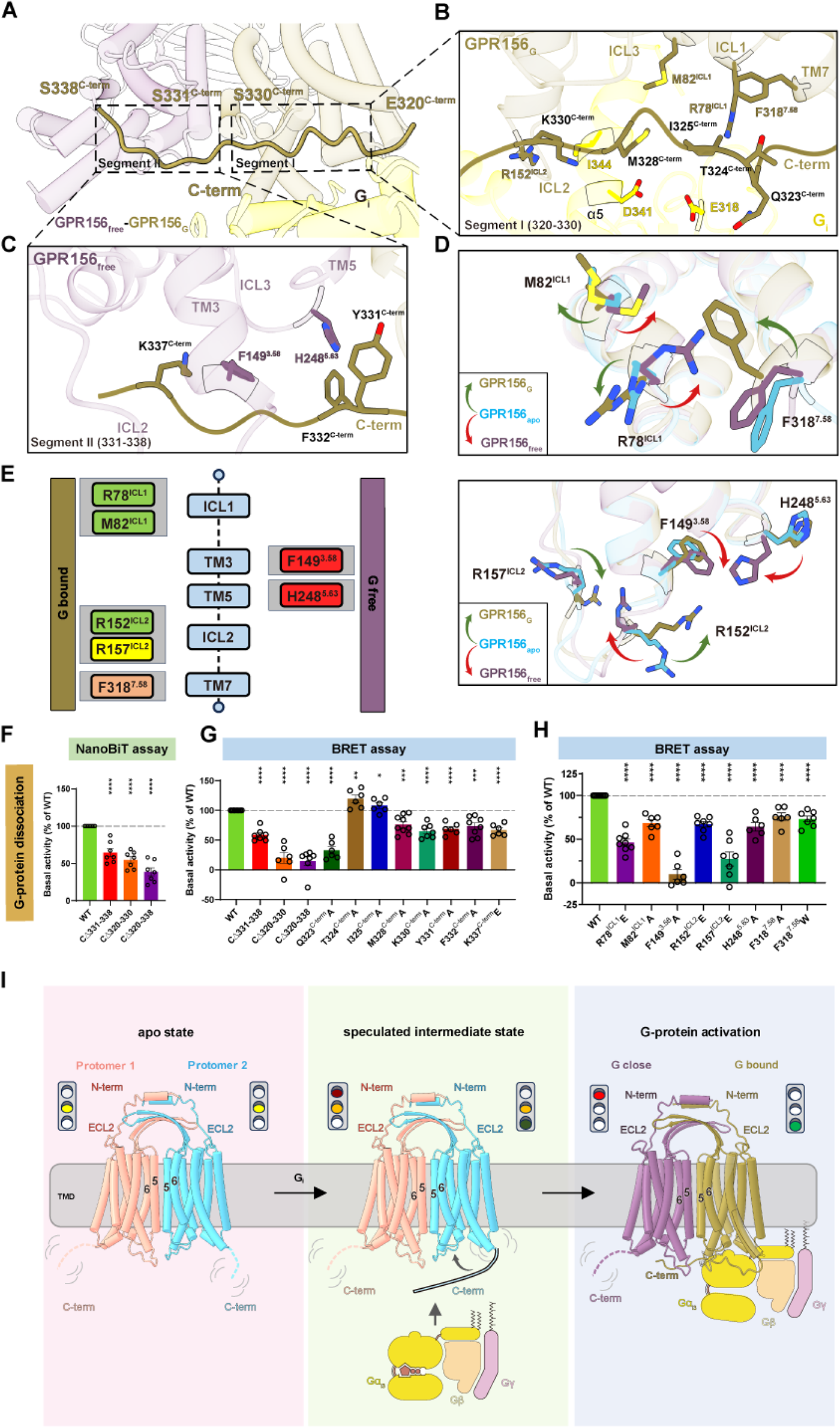
GPR156–Gi coupling and the special role of the C-terminus. **(A)** Close-up views of the GPR156-Gi3 complex at the cytoplasmic end, showing a long C-terminal tail (320–338aa). **(B** and **C)** Detailed interactions of the C-terminal tail (320–330aa) of GPR156G with the TMD of GPR156G and Gαi (B), and detailed interactions of the C-terminal tail (331–338aa) of GPR156G with the TMD of GPR156free (C). **(D)** Close-up views of the seven key residues in the GPR156G and GPR156free subunits show conformational changes upon G protein coupling. **(E)** Schematic representation of the seven key residues’ contacts between the GPR156G and GPR156free subunits and the C-terminal tail of GPR156G and Gαi. Residues from the GPR156G subunit are shown on the left, those of the GPR156free subunit are shown on the right. The C-terminal tail of GPR156G–GPR156free is in red, the C-terminal tail of GPR156G–GPR156free is in orange, the C-terminal tail of GPR156G–Gαi is in yellow, and both C-terminal tails of GPR156G–GPR156G and GPR156G–Gαi are in green boxes. **(F)** The basal activity of WT and the three mutant constructs of the C-terminal tail from GPR156, as measured by NanoBiT-based assay. Fig S8E provides the related surface expression level, and Table S4 provides detailed information. **(G** and **H)** The basal activity of WT and mutant versions of GPR156 (including mutant constructs of the C-terminal tail) (G) and mutant constructs of the seven key residues (H) as measured by BRET-based assay. Data are presented as a percentage of WT activity and are shown as the mean ± SEM (bars) from at least six independent experiments performed in technical triplicate with individual data points shown (dots). \**P* < 0.05, \*\**P* < 0.01, \*\*\**P* < 0.001, \*\*\*\**P* < 0.0001 by two-tailed unpaired t-test compared to WT. Fig S8F and G provides the related surface expression level, and Table S3 provides detailed information. **(I)** Schematic diagrams summarizing the conformational changes of the GPR156 homodimers upon G protein coupling.

### Distinct conformation of the C-terminus in class C GPCRs

The stark deviation observed from the known G protein coupling mechanisms of class C GPCRs warrants an in-depth investigation into how GPR156 responds to G protein coupling. Hence, the C-terminus observed solely in the G-bound subunit caught our attention. The discernible density of the C-terminus from the G-bound subunit includes residues 320–338 and lies approximately parallel to the intracellular end of the GPR156 dimer, whereas it was absent in the apo state (Figs. 2, D and E, 6A, and fig. S5). It spans the G protein-binding pocket in the G-bound subunit and extends to the G-free subunit, thus occupying the bottom of the TMD in the G-free subunit (Fig. 6A). This constitutes the initial identification within class C GPCRs of the G-free subunit’s potential G protein pocket being occupied by the C-terminus (Fig. 6A). The G protein pocket of GPR156 aligns with reports on known class C GPCRs (*19, 32, 33*) by forming a shallow pocket primarily involving ICL1, ICL2, and the C-terminus (fig. S14, A to D, and table S3). The interaction area between the α5 helix of Gα_i3_ and GPR156 is 808 Å^2^ (fig. S14E). Of this, the C-terminus of GPR156 accounts for only 10.6% (86 Å^2^), which is a notable contrast to the C-terminus of mGlu2 or CaSR (fig. S14F). Furthermore, comparing GPR156’s G-bound subunit with mGlu2’s or CaSR’s G-bound subunit (*13, 32, 34*) illustrates that, unlike GPR156, their C-terminus extends downward in close proximity to the G protein without extending into the G protein pocket of the G-free subunit (fig. S14G). These findings suggest that the C-terminus of the G-bound subunit of GPR156 may play a previously unknown role.

### The dual role of the GPR156 C-terminus in G-protein coupling

To further elucidate the specific interaction and role of the C-terminus of the G-bound subunit, it can be divided into two segments (Fig. 6A). Segment I (320–330aa) is involved in interactions with the G-bound subunit itself and the G protein (Fig. 6B), while segment II (331–338aa) participates in interactions with the G-free subunit (Fig. 6C). We hypothesized that Segment I is involved in G protein binding, somewhat similar to the role seen in mGlu2 (*32*). Considering the capability of both subunits to bind G protein, Segment II may prevent the G-free subunit from coupling to the G protein. We found that steric hindrance impeded the possibility of both subunits while simultaneously coupling to the G protein heterotrimer (fig. S15A). It is conceivable that if the potential G protein pocket of the G-free subunit is not occupied by the C-terminus, both subunits would competitively couple with G protein simultaneously, leading to mutual interference. Thus, we posit that Segment II serves the purpose of preventing disturbances from excess G protein binding, thus ensuring the stability of G protein activation. To validate our hypothesis, three constructs (CΔ331–338, CΔ320–330, and CΔ320–338) were constructed (fig. S15B) and the impact of C-terminal deletions on basal activity was assessed (Fig. 6, F to G). In comparison to the WT receptor, the basal activity of all three truncation mutants significantly decreased (Fig. 6, F and G; fig. S8, E and F; and tables S3 and S4), indicating that both segments at the C-terminus have an influence on the constitutive activity of GPR156.

In detail, Q323^C-term^ in Segment I forms potential hydrogen bond with E318^G.h4s6.12^ of the G protein (Fig. 6B). Additionally, M328^C-term^ interacts with D341^G.H5.13^ of the G protein and M82^ICL1^ of the G-bound subunit, while K330^C-term^ engages in interactions with I344^G.H5.16^ of the G protein and R152^ICL2^ of the G-bound subunit (Fig. 6B). Furthermore, T324^C-term^ and I325^C-term^ form interactions with F318^7.58^ and R78^ICL1^ of the G-bound subunit, respectively (Fig. 6B). In Segment II, F332^C-term^ and K337^C-term^ establish interactions with H248^5.63^ and F149^3.58^ of the G-free subunit, respectively, while Y331^C-term^ engages in interactions with H248^5.63^ of the G-free subunit (Fig. 6C). Compared to the WT receptor, mutations of Y331^C-term^, F332^C-term^, and K337^C-term^ in Segment II result in a decrease in basal activity (Fig. 6G; fig. S8F; and table S3). Notably, mutation of Q323^C-term^ in Segment I leads to a more than 50% reduction in basal activity (Fig. 6G), underscoring the crucial role of the C-terminus of the G-bound subunit in the G protein pocket of the G-bound subunit in GPR156 coupling to G_i3_.

After G protein coupling, the side chains of seven amino acids were found to undergo distinct conformational changes in the G protein pocket of G-bound subunit and potential pocket of G-free subunit (Fig. 6, D and E). Mutations of these seven crucial amino acids resulted in varying degrees of reduction in basal activity (Fig. 6H; fig. S8G; and table S3). Notably, F149^3.58^ underwent conformational change only in the G-free pocket and interacts solely with the K337^C-term^ of the G-bound subunit (Fig. 6, C and D). Consistent with this, mutations of F149^3.58^ led to a reduction in basal activity by over half (Fig. 6H), further highlighting the important role of the C-terminus of the G-bound subunit in occupying the potential G protein pocket of the G-free subunit during GPR156–G_i3_ coupling. Combining the aforementioned analysis with functional experiments, we confirm the dual role of the C-terminus of GPR156’s G-bound subunit in the G protein coupling process. On the one hand, the C-terminus promotes G protein coupling in the G protein pocket of the G-bound subunit (320–330aa). On the other hand, the C-terminus occupies the bottom of the TMD of the G-free subunit (331–338aa) so as to impede unwanted perturbations from additional G protein binding.

## Discussion

This study presents the cryo-EM structures of apo GPR156 and the GPR156-G_i3_ complex. Here, we describe a homotypic dimeric form and provide a dynamic view of G protein coupling that is distinct from all known class C GPCRs, and we metaphorically refer to the dual function of the C-terminus as a traffic light (Fig. 6I). Prior to G protein coupling, GPR156 adopts a symmetrical dimeric form, with both subunits exhibiting nearly identical conformation and possessing the ability to couple with G proteins, analogous to being in a yellow light “ready” state. As G protein coupling initiates, there is no rearrangement at the dimeric interface, and the TMDs of the dimer undergo no significant rotation or translation. We hypothesize that the C-terminus of the GPR156 subunit proximal to the G protein interacts with the G protein (e.g., Q323^C-term^–E318^G.h4s6.12^). After G protein coupling, the C-terminus of the G-bound subunit aligns parallel to the intracellular end of the dimer. On the one hand, this facilitates the binding of the G protein of the G-bound subunit, resembling a green light in the open state. On the other hand, it occupies the bottom of the TMD in the G-free subunit, impeding the interference caused by excess G protein binding, akin to a red light in the closed state.

Moreover, the specificity of the small extracellular region of GPR156 is indisputable. The absence of the VFT region, the elongated stalk, and the disulfide bond between ECL2 and TM3 render GPR156 incapable of exerting the conserved functions observed in the activation state of all known class C GPCRs. In addition, there is also a subclass of GPCRs with short N-terminus similar to that of GPR156 in class C GPCRs, including GPRC5A (N-terminus: 26 residues), GPRC5B (N-terminus: 54 residues), GPRC5C (N-terminus: 48 residues), and GPRC5D (N-terminus: 21 residues) (Fig. 2B). A question worth considering is how these class C GPCRs without a large extracellular region form dimers and whether the dimeric form of GPR156 might represent a novel subclass of class C GPCR dimers, and this may be worthy of further study.

An excellent work has recently been reported on GPR156 in the G_o_-free and G_o_-coupled states (*35*), and our physiological and structural analyses presented here have given us further valuable and novel insights into those findings. By comparing the mass spectrometry data with samples of two class A GPCRs known to lack double-chain phospholipids, we have identified the differential enrichment of PG 36:2 in our GPR156 fraction, but not phosphatidylcholine (PC) molecules with corresponding aliphatic chain lengths (fig. S13, A and B). Thus, we speculate that the phospholipids bound within the TMD are more likely to be PG rather than PC. Combining the previous studies of phospholipids in GABA_B_ receptors with the highest homology to GPR156 (*26, 30, 31*), our MD simulation analysis (fig. S13, G and H), and our structural analysis of the toggle switch motif (fig. S13, I to L), we propose that phospholipids within GPR156 act to maintain the integrity and stability of the receptor but might not activate the receptor as recently reported. However, the special dimer interface (TM5/6-TM5/6) and the lack of ionic lock in GPR156 may be the reasons for maintaining the active-like conformation in apo state.

More importantly, our investigation reveals unique features in GPR156’s interaction with the G protein, highlighting a novel binding pattern with G_i3_ that is distinct from the G_o_ coupling that has recently been observed (*35*). Notably, it has been reported that G_o_ is not expressed in auditory HCs, and only G_i2_ and G_i3_ are involved in the orientation effect of GPR156 on HCs (*3*). Therefore, the dual functional role of the C-terminus in the GPR156–G_i3_ complex is more representative for at least the role of GPR156 in auditory function.

Taken together, our investigation reveals the indispensable role of GPR156 in ensuring the normal operation of uninterrupted auditory function across developmental and maturation phases. Our observations further provide structural information regarding the atypical activation mechanism of GPR156, which is characterized by rapid and stable constitutive coupling to G_i3_ protein, making it well-suited for auditory function.

## Materials and methods

### Animals

C57 WT mice were used in all experiments. All experiments were approved by the Institutional Animal Care and Use Committee of Southeast University, China, and all efforts were made to minimize the number of mice used.

### Plasmid design and AAV purification

The GPR156-shRNA sequence was designed with the help of the shRNA-design website (Thermo Fischer Scientific). The GPR156-shRNA was inserted into the AAV vector tagged with the fluorescent protein mNeonGreen. AAV.7m8 was used as the AAV capsid because it can efficiently infect HCs (*36, 37*), and this along with the targeted plasmid and helper plasmid were delivered to HEK-293T cells at molar ratio 1:1:1 by linear polyethylenimine (Yeasen, 40816ES02). AAV purification and titer assays were performed according to the previously published protocol (*38, 39*). The GPR156-shRNA1 sequence was cgg agc atg caa tgt agc ttt.

### AAV injection in mice

The neonatal mice were anesthetized by cooling on ice. The skin of the neck of the left ear was then clipped, and the fat and muscle were pared away to expose the round window membrane. For the neonatal mice, the AAVs were injected into the cochlea via the round window membrane, and the volume was controlled at 1.5 µL.

The adult mice were anesthetized by tribromoethanol (500 mg/kg). The hair was cleared in the left neck area, and a small cut was made in the neck skin to separate the fat and muscle, thus exposing the posterior semicircular canal into which a small hole was drilled. All of the AAVs were injected into the cochlea via the posterior semicircular canal, and the volume was controlled at 2 µL. After the AAV injection, the wound was sutured and sterilized.

### ABR testing

The adult mice were anesthetized by tribromoethanol (500 mg/kg). The closed-field ABR thresholds were measured using tones of different frequencies (4 kHz, 8 kHz, 12 kHz, 16 kHz, and 32 kHz) and different sound intensities (15–90 dB) using a TDT system III workstation (Tucker-Davis Technologies, RZ6).

### Immunofluorescence staining

Samples were fixed with 4% PFA and decalcified with 0.5 M EDTA. The samples were then blocked with 10% donkey serum for 1–2 h and incubated with primary antibodies overnight at 4 °C. The primary antibodies were Myosin7a (Proteus Bioscience, 1:1000 dilution) and Phalloidin (Invitrogen, 1:1000 dilution, A22287). The next day, the secondary antibodies were incubated at room temperature for 1 h. Confocal images were obtained on a confocal microscope (Zeiss LSM 900).

### Expression and purification of apo GPR156

For the expression of apo GPR156, human embryonic kidney (HEK) 293 GnTI^-^ cells were grown in suspension culture at 37 °C in 8% CO_2_ using 293 freestyle media (M293TII, SMM). When the cells had grown to a density of 2.8 × 10^6^ cells/ml, they were infected with recombinant baculoviruses carrying the GPR156 plasmid and further cultured for 18 h at 37 °C. Then 10 mM sodium butyrate was added, and the cells were further incubated for 72 h at 30 °C. Finally, the cells were collected by centrifugation at 1,000 × *g*, washed in PBS, and stored at −80 °C until further use.

The cells were homogenized in a buffer containing 50 mM HEPES, pH 7.5, 150 mM NaCl, 0.025 mM TCEP, 1/100 DNAase, 10% glycerol and a cocktail of protease inhibitors (Roche, Basel, Switzerland). Then the membrane was solubilized for 3 h at 4 °C by adding 0.5% (w/v) lauryl maltoseneopentyl glycol (LMNG, Anatrace), 0.1% (w/v) cholesteryl hemisuccinate (CHS, Anatrace). The cell membrane was then pelleted by ultracentrifugation at 30,000 g for 45min. After that, the supernatant was bound to an MBP column for 2h. The resin was washed by the wash buffer consisting of 50 mM HEPES, pH 7.5, 150 mM NaCl, 0.01% LMNG, 0.002% CHS, 0.025 mM TCEP, 10% glycerol. The protein then was eluted by the same buffer with 10mM Maltose. The eluted GPR156 was subjected to 3C protease cleavage for the removal of 2xMBP tag. Finally, the GPR156 was concentrated in a 100-kDa cutoff Vivaspin (Millipore) filter and subjected to Superose 6 (GE Healthcare) Increase column (GE Healthcare) in 50 mM HEPES, pH 7.5, 150 mM NaCl, 0.002% LMNG and 0.0004% CHS. The peak fractions were collected and concentrated to 9.5 mg/mL for further cryo-EM studies. Then the peak fractions were also verified by western blot and SDS-PAGE analysis, and the main antibodies used were Anti-Strep-tag II primary antibody (Abcam, ab76950) and goat anti-rabbit secondary antibody (Abclonal, AS014).

### Expression and purification of GPR156-G_i3_ complex

The purification steps of GPR156-G_i3_ complex were essentially the same as those for apo GPR156, except for incubating the concentrated sample with 1.3 moles of scFv16 (which was expressed and purified as previously described (*19*)) at 4°C for 1 h before applying the Superose6 Increase column.

### Cryo-EM sample preparation and data acquisition

For cryo-EM grid preparation, three microlitres of the purified apo GPR156, GPR156-G_i3_ complex (apo: 9.5 mg/ml, G_i3_-bound: 4.5 mg/ml) was applied onto the glow-discharged holey carbon grids (Quantifoil, R1.2/1.3, 300 mesh). The grids were blotted for 3.0 s with a blot force of 10 at 4 ℃, 100% humidity, and then plunge-frozen in liquid ethane using Vitrobot Mark IV (Thermo Fischer Scientific). Cryo-EM data collection was performed on Titan Krios at 300 kV accelerating voltage at the Core Facilities, Liangzhu laboratory, Zhejiang University equipped with CFEG and the Center of Cryo-Electron Microscopy, Zhejiang University equipped with XFEG. Both instruments were equipped with Selectris energy filter, and Falcon 4 direct electron detector. EPU software was used for automated data collection according to standard procedures. Magnification of ×130,000 was used for imaging, yielding a pixel size of 0.93 Å on images. The defocus range was set from −1.0 to −2.5 μm. Each micrograph was dose-fractionated to 40 frames under a dose rate of about 8.7 electrons per pixel per second, with a total exposure time of 6 s, resulting in a total dose of about 52 electrons per Å^2^.

### Image processing and 3D reconstruction

Cryo-EM image stacks were aligned using RELION 4.0 (*40*). Contrast transfer function (CTF) parameters were estimated by Gctf v1.18 (*41*). The following data processing was performed using RELION 4.0 and CryoSPARC v4.4.0 (*42*).

For apo GPR156, automated particle selection using Topaz picking in RELION produced 17,010,572 particles. The particles were imported to CryoSPARC for several rounds of 2D classification and ab-initio reconstruction to generate the initial reference maps, followed by several rounds of heterogeneous refinement in CryoSPARC. The good particles were subjected to Non-uniform Refinement and generated a map with an indicated global resolution of 3.09 Å in CryoSPARC, which were sharpened with deepEMhancer. This map was used for subsequent model building and analysis.

For GPR156-Gi_3_ complex, 12,058,641 particles produced from the automated particle picking were subjected to 2D classification, ab-initio reconstruction and 3D heterogeneous refinement in CryoSPARC, resulting in 1,336,840 well-defined particles. Further several rounds of 3D classification generating one high-quality subset. The final good particles were subjected to a final round of 3D refinement, CTF refinement and Bayesian polishing, generating a map with an indicated global resolution of 2.40 Å at a Fourier shell correlation of 0.143. The final map was sharpened with deepEMhancer and used for subsequent model building and analysis.

### Model building and refinement

The AlphaFold2 predicted structure was used to generate the initial model of GPR156 (*43*). The atomic coordinates of G_i3_ protein from the structure of the mGlu4–G_i3_ complex (PDB: 7E9H) were used to generate the initial model of the G_i3_ protein (*44*). Models were manually docked into the density maps of apo GPR156 and GPR156-G_i3_ complex using UCSF Chimera. Then, the initial models were subjected to flexible fitting using Rosetta and were further rebuilt in Coot and real-space-refined in Phenix (*45–47*). The final refinement statistics were validated using the module ‘comprehensive validation (cryo-EM)’ in Phenix. The refinement statistics are provided in table S2. Structural figures were created using UCSF Chimera X package (*48*).

### Identification of phospholipid ligands by LC-MS/MS

The sample preparation of the phospholipid specifically bound to GPR156 was performed following previous methods (*49*). The samples of GPR156 were prepared in three independent replicates. Additionally, two well-established class A GPCRs (including GPR34 (*50, 51*) and GPR174 (*51–53*)), each with two replicates, were employed as control groups where no phospholipids with two chains were present. A Dionex U3000 UHPLC (Waters Corporation, Milford, USA) fitted with Q-Exactive mass spectrometer equipped with heated electrospray ionization (ESI) source (Thermo Fisher Scientific, Waltham, MA, USA) was used to analyze the metabolic profiling in both ESI positive and ESI negative ion modes. An ACQUITY UPLC BEH C8 column (1.7 μm, 2.1 × 100 mm) were employed in both positive and negative modes. The binary gradient elution system consisted of (A) acetonitrile: water (60:40, v:v, containing 10 mmol/L ammonium formate) and (B) acetonitrile: isopropanol (10:90, v:v, containing 10 mmol/L ammonium formate) and separation was achieved using the following gradient:0 min, 30% B; 3 min, 30% B; 5 min, 62% B; 15 min, 82% B; 16.5 min, 99% B; 18 min, 99% B;18.1 min, 30% B, 22 min, 30% B. The flow rate was 0.26 mL/min and column temperature was 55°C. Positive: Spray Voltage (kV): +3.5; Capillary temperature 300°C; Aux gas heater 350 °C; Sheath gas flow rate (Arb): 45; Aux gas flow rate (Arb): 10; S-lens RF level: 50; Mass range (m/z): 150-1500, Full ms resolution: 70000; MS/MS resolution:17500; TopN: 10; NCE/stepped NCE: 25, 35, 45. Negative: Spray Voltage (kV): -3.0; Capillary temperature 300 °C; Aux gas heater temperature 350 °C; Sheath gas flow rate (Arb): 45; Aux gas flow rate (Arb): 10; S-lens RF level: 50; Mass range (m/z): 150-1500, Full ms resolution: 70000; MS/MS resolution:17500; TopN: 10; NCE/stepped NCE: 25, 35, 45.

The peak extraction and lipids identification were acquired using MSDIAL (v.4.9). Candidate phospholipid molecules are annotated by the following parameters: MS/MS assigned = TRUE; Total score ≥ 70 (Total score is based on accurate mass, isotope ratio, retention time, and MS/MS similarities); can be detected in all three independent replicates of GPR156; log2 fold change (logFC) ≥ 2. The identified endogenous phospholipid bound to GPR156 was phosphatidylglycerol (PG) 36:2 (PG (18:1_18:1); International Chemical Identifier (InChI) key: DSNRWDQKZIEDDB-UHFFFAOYSA-N), which MS2 spectra Precursor: Reference Mass is 773.53375: 773.53381.

### Molecular dynamics simulations

MD simulations were performed essentially as previously described (*31*) with modifications. From the structure of the GPR156 dimer complex, the two transmembrane helix core-bound phospholipids were directly removed, and the retained GPR156-dimer complex was used as the initial configuration for our MD simulations. The GPR156-dimer complex was embedded into a membrane made of 3:1 POPC:POPE and a cholesterol bilayer, which was built using the Membrane Builder module on the CHARMM-GUI server (*54*). The resulting system had a total of 240 lipid and 7 cholesterol molecules and was solvated in a cubic box with TIP3P waters and 0.15 M Na^+^/Cl^−^ ions. The CHARMM36m forcefield was used to describe the system (*55*), and all MD simulations were performed using GROMACS-2019.4 (*56*). After 5000 steps of energy minimization performed by the steepest descent algorithm, a 250 ps NVT equilibration simulation was performed at 310 K. Subsequently, a cumulatively 1.5 ns NPT equilibration to 1 atm was performed using the Berendsen barostat (*57*). Long-range electrostatic interactions were treated with the particle-mesh Ewald method (*58, 59*). The short-range electrostatic and van der Waals interactions both used a cutoff of 10 Å. All bonds were constrained by the LINCS algorithm (*60*). Finally, to escape the uncertainties of the sampling results, three replicates of the production run with different initial velocities were performed. The cavity volume of the transmembrane helices core was calculated with Epock (1.0.5) (*61*) in VMD (*62*).

### Enzyme-linked immunosorbent (ELISA) assay

The cell-surface expression of WT GPR156 or its mutants was detected using an ELISA assay as previously described (*19*). In brief, HEK293T cells were transfected with plasmids of a mixture of WT GPR156 or mutants, Gα_i3_–LgBiT, Gγ–SmBiT, and Gβ_1_ using Lipofectamine 3000 (Thermo Fisher Scientific) in 500 μl of Opti-MEM (Thermo Fisher Scientific). Twenty-four hours post transfection, cells were distributed into 96-well plates with a white non-transparent bottom and further incubated for 24 hours at 37 °C. The HEK293T cells were then fixed with 4% paraformaldehyde and blocked with 1% bovine serum albumin (BSA) for 1 hour. Luminescence detection was performed using SuperSignal ELISA Femto Maximum Sensitivity substrate (ThermoFisher Scientific) after binding of antibodies coupled to horse-radish peroxidase, and luminescence was measured with a Multimode Microplate Reader (Tecan).

Adjusting the transfection levels of both the WT GPR156 and its mutants was essential to ensuring that the mutants exhibited cell surface expression comparable to WT without significant statistical differences. This adjustment facilitated a meaningful comparison in the G_i_ dissociation assay.

### Bioluminescence resonance energy transfer (BRET)-G-protein dissociation assay

G_i_ activation was assessed by the BRET dissociation assay measuring the proximal interaction between the α and γ subunits of the G_i_ protein. The G protein BRET probes, including Gα_i3_-Rluc8, Gβ_1_, and Gγ_2_-Venus, were generated according to a previous publication (*30*). HEK293T cells were transfected with WT GPR156 or mutants and G_i_ protein probes. Twenty-four hours after transfection, the cells were divided into 96-well plates and incubated for additional 24 hours at 37 °C. The BRET signal was quantified following the introduction of the luciferase substrate coelenterazine h (10 μM) employing a Multimode Microplate Reader (Tecan) equipped with BRET filter sets. The BRET signal was determined as the ratio of light emission at 520 nm/460 nm.

### NanoBiT-G-protein dissociation assay

G_i_ activation was also assessed by the NanoBiT-based dissociation assay measuring the proximal interaction between the α and γ subunits of the G_i_ protein. The transfection system was the same as that in the ELISA assay described above. G protein NanoBiT probes, including Gα_i3_-LgBiT, Gβ_1_, and Gγ_2_-SmBiT, were generated according to a previous publication (*19*). After 1 day of transfection, cells were divided into 96-well plates and incubated for an additional 1 day at 37 °C. The NanoBiT signal was measured using a Multimode Microplate Reader (Tecan).

### Statistical analysis

All data are presented as the mean ± standard error of the mean (SEM), and statistical analyses were performed using GraphPad Prism 9 software. Bar graphs depict the differences of each mutant relative to the WT receptor. Data were from at least six independent experiments, each conducted in triplicate. \**P* < 0.05, \*\**P* < 0.01, \*\*\**P* < 0.001, \*\*\*\**P* < 0.0001 (two-tailed unpaired t test compared to WT receptor).

### Data availability

All data generated in this study are included in the main text or Supplementary Information. The cryo-EM density maps and the atomic coordinates have been deposited in the Electron Microscopy Data Bank (EMDB) and Protein Data Bank (PDB) databases under accession codes EMD-39345 and 8YJP for apo-GPR156; and EMD-39356 and 8YK0 for the GPR156–G_i3_ complex. Materials are available from the corresponding authors upon request.

## Competing interests

The authors declare no competing interests.

## Acknowledgements

The cryo-EM data were collected at the Cryo-Electron Microscopy Center of Liangzhu laboratory. Cryo-EM specimens were screened at the Center of Cryo-Electron Microscopy, Zhejiang University. Protein purification was performed at the Protein Facilities of Zhejiang University School of Medicine. We thank Shanghai OE Biotech for providing mass spectrometry detection. This project was supported by the National Key Research and Development Program of China (2021YFA1101300 (R.C.), 2021YFA1101800 (R.C.), 2020YFA0113600 (J.Q.), and 2020YFA0112503 (R.C.)); the STI2030-Major Projects (2022ZD0205400 (R.C.)); the National Natural Science Foundation of China (82030029 (R.C.), 92149304 (R.C.), 82371162 (R.C.), 82371161 (R.C.), 92353303 (Y.Z.), 32141004 (Y.Z.), 32300598 (C.S.), and 82000984 (J.Q.)); the Ministry of Science and Technology (2019YFA050880 (Y.Z.)); the Key R&D Projects of Zhejiang Province (2021C03039 (Y.Z.)); the Leading Innovative and Entrepreneur Team Introduction Program of Zhejiang (2020R01006 (Y.Z.)); the “Pioneer” and “Leading Goose” R&D Program of Zhejiang (2024C03147 (Y.Z.)); the China National Postdoctoral Program for Innovative Talents (BX20200082 (J.Q.)); the China Postdoctoral Science Foundation (2023TQ0056 (M.L.), 2023M730575 (M.L.), GZC20230435 (M.L.), and 2020M681468 (J.Q.)); the Science and Technology Department of Sichuan Province (2021YFS0371 (R.C.)); the Shenzhen Science and Technology Program (JCYJ20210324125608022 (R.C.) and JCYJ20190814093401920 (R.C.)); the Open Research Fund of State Key Laboratory of Genetic Engineering, Fudan University (SKLGE-2104 (R.C.)); the 2022 Open Project Fund of Guangdong Academy of Medical Sciences to P.N.W. (YKY-KF202201(R.C.)); and the Fundamental Research Funds for the Central Universities (R.C. and Y.Z.).

## Author contributions

R.C. and Y.Z. conceived and supervised the whole project; L.Z. and X.M. designed the AAV strategy; L.Z. performed the animal experiments; X.M. and J.G. designed the constructs and expressed and purified apo GPR156 and the GPR156–G_i3_ complex; D.-D.S. evaluated the sample by negative-stain EM; L.-N.C. prepared the cryo-EM grids; L.-N.C., S.-K.Z., J.Q., and S.-Y.J. collected the cryo-EM data; L.-N.C. and C.M. performed cryo-EM map calculation and model building; M.L., X.M., and J.G. generated the mutant constructs; M.L. performed the *in vitro* cellular functional assays; X.M. and Y.-J.D. prepared the samples for the mass spectrometry studies; P.C. and Y.L. performed bioinformatics analysis of the mass spectrometry data; K.X. performed the MD simulations; X.M. and L.-N.C. performed the structural analysis; X.M. and L.-N.C. prepared the figures; C.S., M.L., L.Z., J.G., Q.S., H.Z., J.Q., L.M., and J.L. participated in data analysis; X.M. and L.-N.C. prepared the draft of the manuscript; R.C., Y.Z., C.M., and J.L. wrote the manuscript with inputs from all the authors.

## Notes

### Competing Interest Statement

The authors have declared no competing interest.

## References

1. K. S. Kindt et al., EMX2-GPR156-Galphai reverses hair cell orientation in mechanosensory epithelia. Nat Commun 12, 2861 (2021).doi:10.1038/s41467-021-22997-1.

2. D. Greene et al., Genetic association analysis of 77,539 genomes reveals rare disease etiologies. Nat Med, (2023).doi:10.1038/s41591-023-02211-z.

3. A. Jarysta et al., Inhibitory G proteins play multiple roles to polarize sensory hair cell morphogenesis. eLife 12, (2023).doi:10.7554/eLife.88186.1.

4. K. J. Charles, A. R. Calver, S. Jourdain, M. N. Pangalos, Distribution of a GABAB-like receptor protein in the rat central nervous system. Brain Res 989, 135–146 (2003).doi:10.1016/s0006-8993(03)03163-9.

5. L. Chun, W. H. Zhang, J. F. Liu, Structure and ligand recognition of class C GPCRs. Acta Pharmacol Sin 33, 312–323 (2012).doi:10.1038/aps.2011.186.

6. T. Laboute et al., Orphan receptor GPR158 serves as a metabotropic glycine receptor: mGlyR. Science 379, 1352–1358 (2023).doi:10.1126/science.add7150.

7. D. N. Patil et al., Structure of the photoreceptor synaptic assembly of the extracellular matrix protein pikachurin with the orphan receptor GPR179. Sci Signal 16, eadd9539 (2023).doi:10.1126/scisignal.add9539.

8. B. Bettler, K. Kaupmann, J. Mosbacher, M. Gassmann, Molecular structure and physiological functions of GABA(B) receptors. Physiol Rev 84, 835–867 (2004).doi:10.1152/physrev.00036.2003.

9. A. B. Caniceiro, B. Bueschbell, A. C. Schiedel, I. S. Moreira, Class A and C GPCR Dimers in Neurodegenerative Diseases. Curr Neuropharmacol 20, 2081–2141 (2022).doi:10.2174/1570159X20666220327221830.

10. J. Kniazeff, L. Prezeau, P. Rondard, J. P. Pin, C. Goudet, Dimers and beyond: The functional puzzles of class C GPCRs. Pharmacol Ther 130, 9–25 (2011).doi:10.1016/j.pharmthera.2011.01.006.

11. H. Wu et al., Structure of a class C GPCR metabotropic glutamate receptor 1 bound to an allosteric modulator. Science 344, 58–64 (2014).doi:10.1126/science.1249489.

12. J. Du et al., Structures of human mGlu2 and mGlu7 homo- and heterodimers. Nature 594, 589–593 (2021).doi:10.1038/s41586-021-03641-w.

13. X. Wang et al., Structural insights into dimerization and activation of the mGlu2-mGlu3 and mGlu2-mGlu4 heterodimers. Cell Res 33, 762–774 (2023).doi:10.1038/s41422-023-00830-2.

14. T. K. Attwood, J. B. Findlay, Fingerprinting G-protein-coupled receptors. Protein Eng 7, 195–203 (1994).doi:10.1093/protein/7.2.195.

15. A. Koehl et al., Structural insights into the activation of metabotropic glutamate receptors. Nature 566, 79–84 (2019).doi:10.1038/s41586-019-0881-4.

16. A. Ellaithy, J. Gonzalez-Maeso, D. A. Logothetis, J. Levitz, Structural and Biophysical Mechanisms of Class C G Protein-Coupled Receptor Function. Trends Biochem Sci 45, 1049–1064 (2020).doi:10.1016/j.tibs.2020.07.008.

17. L. R. Watkins, C. Orlandi, In vitro profiling of orphan G protein coupled receptor (GPCR) constitutive activity. Br J Pharmacol 178, 2963–2975 (2021).doi:10.1111/bph.15468.

18. A. R. Calver et al., Molecular cloning and characterisation of a novel GABAB-related G-protein coupled receptor. Brain Res Mol Brain Res 110, 305–317 (2003).doi:10.1016/s0169-328x(02)00662-9.

19. C. Shen et al., Structural basis of GABA(B) receptor-G(i) protein coupling. Nature 594, 594–598 (2021).doi:10.1038/s41586-021-03507-1.

20. A. Koehl et al., Structural insights into the activation of metabotropic glutamate receptors. Nature 566, 79-+ (2019).doi:10.1038/s41586-019-0881-4.

21. Y. Gao et al., Asymmetric activation of the calcium-sensing receptor homodimer. Nature 595, 455-+ (2021).doi:10.1038/s41586-021-03691-0.

22. L. D. Liberman, M. C. Liberman, Postnatal maturation of auditory-nerve heterogeneity, as seen in spatial gradients of synapse morphology in the inner hair cell area. Hear Res 339, 12–22 (2016).doi:10.1016/j.heares.2016.06.002.

23. D. O. Mikaelian, R. J. Ruben, Hearing Degeneration in Shaker-1 Mouse. Correlation of Physiological Observations with Behavioral Responses and with Cochlear Anatomy. Arch Otolaryngol 80, 418–430 (1964).doi:10.1001/archotol.1964.00750040430011.

24. U. H. Isu, S. A. Badiee, E. Khodadadi, M. Moradi, Cholesterol in Class C GPCRs: Role, Relevance, and Localization. Membranes (Basel) 13, (2023).doi:10.3390/membranes13030301.

25. D. N. Patil et al., Cryo-EM structure of human GPR158 receptor coupled to the RGS7-Gbeta5 signaling complex. Science 375, 86–91 (2022).doi:10.1126/science.abl4732.

26. J. Park et al., Structure of human GABA(B) receptor in an inactive state. Nature 584, 304–309 (2020).doi:10.1038/s41586-020-2452-0.

27. H. Shaye et al., Structural basis of the activation of a metabotropic GABA receptor. Nature 584, 298–303 (2020).doi:10.1038/s41586-020-2408-4.

28. E. Jeong, Y. Kim, J. Jeong, Y. Cho, Structure of the class C orphan GPCR GPR158 in complex with RGS7-Gbeta5. Nat Commun 12, 6805 (2021).doi:10.1038/s41467-021-27147-1.

29. A. S. Dore et al., Structure of class C GPCR metabotropic glutamate receptor 5 transmembrane domain. Nature 511, 557–562 (2014).doi:10.1038/nature13396.

30. C. Mao et al., Cryo-EM structures of inactive and active GABA(B) receptor. Cell Res 30, 564–573 (2020).doi:10.1038/s41422-020-0350-5.

31. M. M. Papasergi-Scott et al., Structures of metabotropic GABA(B) receptor. Nature 584, 310–314 (2020).doi:10.1038/s41586-020-2469-4.

32. A. B. Seven et al., G-protein activation by a metabotropic glutamate receptor. Nature 595, 450–454 (2021).doi:10.1038/s41586-021-03680-3.

33. F. He et al., Allosteric modulation and G-protein selectivity of the Ca(2+)-sensing receptor. Nature 626, 1141–1148 (2024).doi:10.1038/s41586-024-07055-2.

34. F. He et al., Allosteric modulation and G-protein selectivity of the Ca(2+)-sensing receptor. Nature, (2024).doi:10.1038/s41586-024-07055-2.

35. J. Shin et al., Constitutive activation mechanism of a class C GPCR. Nat Struct Mol Biol, (2024).doi:10.1038/s41594-024-01224-7.

36. K. Isgrig et al., AAV2.7m8 is a powerful viral vector for inner ear gene therapy. Nat Commun 10, 427 (2019).doi:10.1038/s41467-018-08243-1.

37. J. Zhu et al., Refining surgical techniques for efficient posterior semicircular canal gene delivery in the adult mammalian inner ear with minimal hearing loss. Sci Rep 11, 18856 (2021).doi:10.1038/s41598-021-98412-y.

38. F. Tan et al., AAV-ie enables safe and efficient gene transfer to inner ear cells. Nat Commun 10, 3733 (2019).doi:10.1038/s41467-019-11687-8.

39. L. Zhang et al., AAV-Net1 facilitates the trans-differentiation of supporting cells into hair cells in the murine cochlea. Cell Mol Life Sci 80, 86 (2023).doi:10.1007/s00018-023-04743-6.

40. S. H. Scheres, Processing of Structurally Heterogeneous Cryo-EM Data in RELION. Methods Enzymol 579, 125–157 (2016).doi:10.1016/bs.mie.2016.04.012.

41. K. Zhang, Gctf: Real-time CTF determination and correction. J Struct Biol 193, 1–12 (2016).doi:10.1016/j.jsb.2015.11.003.

42. A. Punjani, J. L. Rubinstein, D. J. Fleet, M. A. Brubaker, cryoSPARC: algorithms for rapid unsupervised cryo-EM structure determination. Nat Methods 14, 290–296 (2017).doi:10.1038/nmeth.4169.

43. J. Jumper et al., Highly accurate protein structure prediction with AlphaFold. Nature 596, 583–589 (2021).doi:10.1038/s41586-021-03819-2.

44. S. Lin et al., Structures of G(i)-bound metabotropic glutamate receptors mGlu2 and mGlu4. Nature 594, 583–588 (2021).doi:10.1038/s41586-021-03495-2.

45. R. Y. Wang et al., Automated structure refinement of macromolecular assemblies from cryo-EM maps using Rosetta. Elife 5, (2016).doi:10.7554/eLife.17219.

46. P. D. Adams et al., PHENIX: a comprehensive Python-based system for macromolecular structure solution. Acta Crystallogr D Biol Crystallogr 66, 213–221 (2010).doi:10.1107/S0907444909052925.

47. P. Emsley, K. Cowtan, Coot: model-building tools for molecular graphics. Acta Crystallogr D Biol Crystallogr 60, 2126–2132 (2004).doi:10.1107/S0907444904019158.

48. T. D. Goddard et al., UCSF ChimeraX: Meeting modern challenges in visualization and analysis. Protein Sci 27, 14–25 (2018).doi:10.1002/pro.3235.

49. X. Qu et al., Structural basis of tethered agonism of the adhesion GPCRs ADGRD1 and ADGRF1. Nature 604, 779–785 (2022).doi:10.1038/s41586-022-04580-w.

50. A. Xia et al., Cryo-EM structures of human GPR34 enable the identification of selective antagonists. Proc Natl Acad Sci U S A 120, e2308435120 (2023).doi:10.1073/pnas.2308435120.

51. G. Liu, X. Li, Y. Wang, X. Zhang, W. Gong, Structural basis for ligand recognition and signaling of the lysophosphatidylserine receptors GPR34 and GPR174. PLoS Biol 21, e3002387 (2023).doi:10.1371/journal.pbio.3002387.

52. J. Liang et al., Structural basis of lysophosphatidylserine receptor GPR174 ligand recognition and activation. Nat Commun 14, 1012 (2023).doi:10.1038/s41467-023-36575-0.

53. Y. Nie et al., Specific binding of GPR174 by endogenous lysophosphatidylserine leads to high constitutive G(s) signaling. Nat Commun 14, 5901 (2023).doi:10.1038/s41467-023-41654-3.

54. S. Jo, T. Kim, V. G. Iyer, W. Im, CHARMM-GUI: a web-based graphical user interface for CHARMM. J Comput Chem 29, 1859–1865 (2008).doi:10.1002/jcc.20945.

55. J. Huang, A. D. MacKerell, Jr., CHARMM36 all-atom additive protein force field: validation based on comparison to NMR data. J Comput Chem 34, 2135–2145 (2013).doi:10.1002/jcc.23354.

56. S. Pronk et al., GROMACS 4.5: a high-throughput and highly parallel open source molecular simulation toolkit. Bioinformatics 29, 845–854 (2013).doi:10.1093/bioinformatics/btt055.

57. H. J. C. Berendsen, J. P. M. Postma, W. F. Vangunsteren, A. Dinola, J. R. Haak, Molecular-Dynamics with Coupling To an External Bath. J Chem Phys 81, 3684–3690 (1984).doi:Doi 10.1063/1.448118.

58. U. Essmann et al., A Smooth Particle Mesh Ewald Method. J Chem Phys 103, 8577–8593 (1995).doi:Doi 10.1063/1.470117.

59. T. Darden, D. York, L. Pedersen, Particle Mesh Ewald - an N.Log(N) Method for Ewald Sums in Large Systems. J Chem Phys 98, 10089–10092 (1993).doi:Doi 10.1063/1.464397.

60. B. Hess, H. Bekker, H. J. C. Berendsen, J. G. E. M. Fraaije, LINCS: A linear constraint solver for molecular simulations. Journal of Computational Chemistry 18, 1463–1472 (1997).doi:Doi 10.1002/(Sici)1096-987x(199709)18:12<1463::Aid-Jcc4>3.3.Co;2-L.

61. B. Laurent et al., Epock: rapid analysis of protein pocket dynamics. Bioinformatics 31, 1478–1480 (2015).doi:10.1093/bioinformatics/btu822.

62. W. Humphrey, A. Dalke, K. Schulten, VMD: Visual molecular dynamics. J Mol Graph Model 14, 33–38 (1996).doi:Doi 10.1016/0263-7855(96)00018-5.

